# Structural basis for the delivery of activated sialic acid into Golgi for sialyation

**DOI:** 10.1101/580449

**Authors:** Emmanuel Nji, Ashutosh Gulati, Abdul Aziz Qureshi, Mathieu Coincon, David Drew

**Author notes:** These authors contributed equally (Emmanuel Nji, Ashutosh Gulati, Abdul Aziz Qureshi).

## Abstract

The decoration of secretory glycoproteins and glycolipids with sialic acid is critical to many physiological and pathological processes. Sialyation is dependent on a continuous supply of sialic acid into Golgi organelles in the form of CMP-sialic acid. Translocation of CMP-sialic acid into Golgi is carried out by the CMP-sialic acid transporter (CST). Mutations in human CST are linked to glycosylation disorders, and CST is important for glycopathway engineering, as it is critical for sialyation efficiency of therapeutic glycoproteins. The mechanism of how CMP-sialic acid is recognized and translocated across Golgi membranes in exchange for CMP is poorly understood. Here we have determined the crystal structure of a eukaryotic CMP-sialic acid transporter in complex with CMP. We conclude that the specificity of CST for CMP-sialic acid is established by the nucleotide CMP to such an extent, they are uniquely able to work both as passive and as (secondary) active antiporters.

## Introduction

Sialic acids decorate the outermost ends of secreted and cell surface expressed glycoproteins and glycolipids ^1,2^. Terminal sialic acid linkages are vital for cellular recognition, immunology and communication ^3^. The cell-surface display of sialic acid is also a receptor for interactions with viruses, such as Influenza and HIV ^1,4^. Sialic acid levels are also increased in cancer cells ^5,6^, which has led to the definition of sialic acids as potential therapeutic targets ^7^. In the nucleus, sialic acid is activated to CMP-sialic acid, where it is delivered into the Golgi lumen by the action of the CMP-Sialic acid transporter (CST) ^8,9^ (Fig. 1a). CSTs belongs to the Solute Carrier SLC35 family of Nucleotide Sugar Transporters (NST), which in humans translocate seven other essential sugars into the ER and/or Golgi for glycosylation in the form of GDP-fucose, UDP-galactose, UDP-glucose, UDP-xylose, UPD-GlcNAc, UDP-GalNAc and UDP-GlcA ^8,10^. SLC35 activities are important to human physiology and have been associated with diseases such as obesity ^11,12^. SLC35 members can influence the efficacy of anti-cancer drugs ^13^, and also transport UDP-glucuronic acid into the liver ER for glucuronidation ^14^, which is an essential process for the metabolism of many drugs, e.g., analgesics, nonsteroidal anti-inflammatory agents, antipsychotics, antivirals, and benzodiazepines ^15^.

**Fig. 1.**
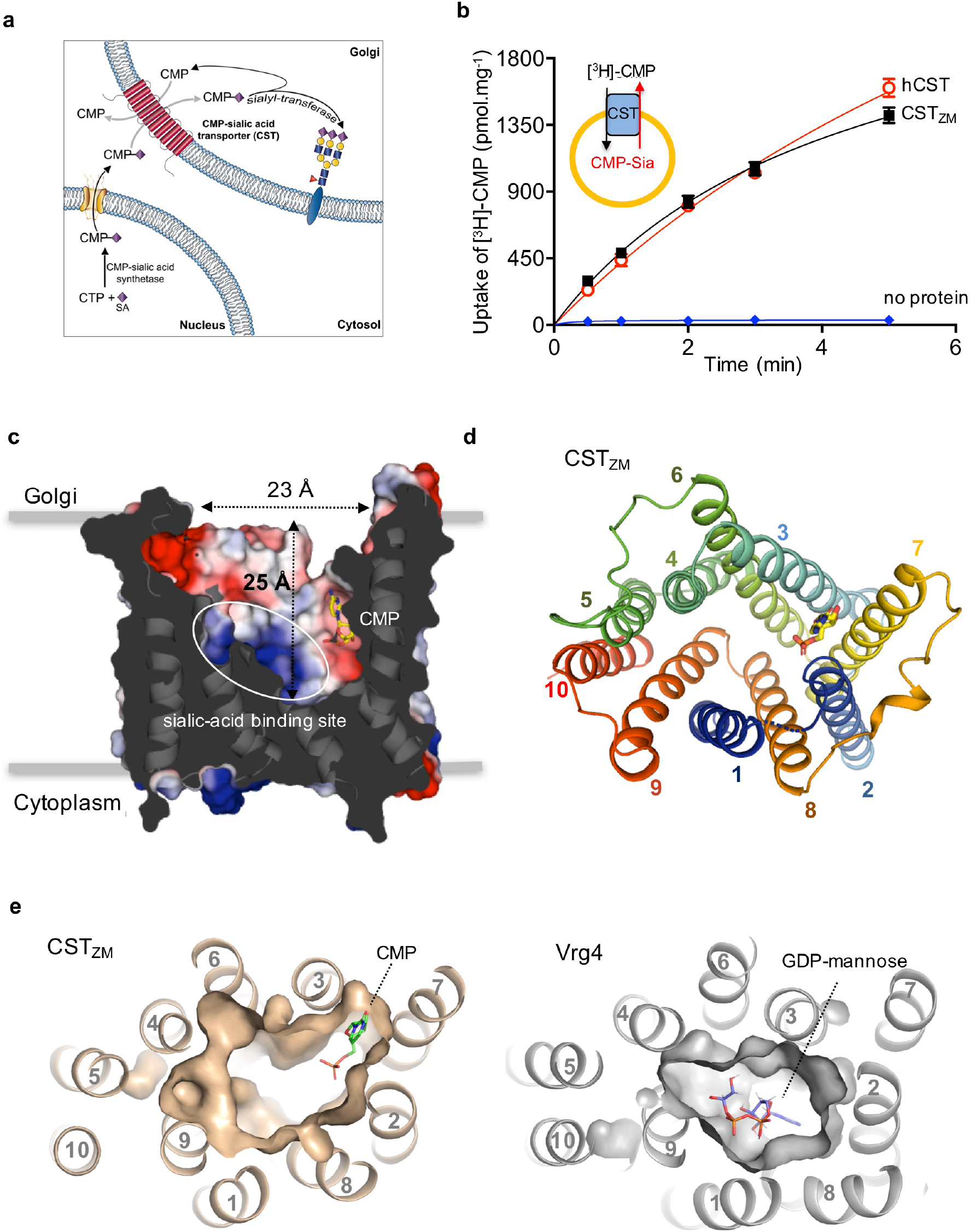
Architecture of CST_ZM_. **a**, Schematic showing the pathway for sialyation of secretory proteins with sialic acid (SA). **b**, Time-dependent uptake of [^3^H]-CMP by CST_ZM_ (black-filled squares) and hCST (red-open circles) in proteoliposomes incubated with CMP-sialic acid. The proteoliposome setup is shown as a schematic in the upper-left hand corner. Non-specific uptake was estimated from intensities derived from liposomes incubated without protein (blue-filled rhombus). Errors bars represent s.e.m. of 3 independent experiments. **c**, Slab through the CST_ZM_, with solvent exposed surface (colored according to the calculated electrostatic potential, presented in blue-positive to red-negative) as viewed within the plane of membrane with CMP (shown as yellow sticks) and the proposed site for sialic acid binding (indicated by the dotted ellipse). **d**, Ribbon representation of the outward-facing CST_ZM_, viewed from the top (TMs shown as cartoon colored as rainbow from blue to red and CMP as yellow sticks). The loop residues 27 to 30 that could not be built are illustrated by a dotted line. **e**, Slab through the surface of outward-facing CST_ZM_ (left panel; light brown) and outward-facing Vrg4 (PDB id: 5OGK right; grey), as viewed from the top, which highlights the difference in binding position of the substrates CMP (left panel; green sticks) and GDP-mannose (right panel; purple sticks) relative to the cavity. Notably, the predicted CMP binding site motif “NIQM” based on structure of Vrg4 ^23^ is located below the surface of the outward-facing CST_ZM_ cavity structure shown in Fig. **1c**.

A number of NST show overlapping substrate specificities ^8,16^, transporting the charged nucleotide-diphosphate sugar (G/UDP-sugar) in exchange for their counter nucleotide monophosphate, G/UMP. CMP-sialic acid transporters are atypical nucleotide-sugar transporters, as they are only found in Golgi organelles and transport sialic acid conjugated to a nucleotide monophosphate *i.e*., CMP-sialic acid is exchanged for CMP ^8,9,17^. Several CST mutations have been linked to human pathologies ^18^ and congenital glycosylation disorders ^19^. Since sialyation efficiency affects the therapeutic efficacy of many therapeutic glycoproteins ^20,21^, CST is also a promising candidate for glycoengineering, as it is the rate limiting factor in the sialyation of glycoproteins ^22^. Recently, an outward-facing structure of a nucleotide-sugar transporter for the *yeast* GDP-mannose transporter (Vrg4) was reported ^23^. However, GDP-mannose is only transported into either the ER or Golgi in non-mammalian organisms such as yeast ^24^. Because of the large evolutionary gap between the GDP-mannose and CMP-sialic acid transporters ^25^, the molecular basis for how activated sialic acid is transported into Golgi is still yet to be understood. Indeed, the poor structural and mechanistic understanding of SLC35 mediated-transport in general, has hampered elucidating their role in human physiology and their targeting in drug development.

## Results

### Crystal structures of the human CMP-Sialic acid transporter homologue CST_ZM_

The CST homologue from *Zea mays* (CST_ZM_) with 28 % sequence identity to human, and ~80% sequence identity to an *Arabidopsis thaliana* CST capable of complementing CMP-sialic acid transport in the model *Chinese Hamster Ovary* cell line Lec2 ^26^, was identified by fluorescent-based screening methods ^27,28^ as a suitable candidate for structural studies (Supplementary Fig. 1a). Consistently, binding studies using the GFP-Thermal Shift (GFP-TS) assay had shown that CST_ZM_ was only significantly thermostabilized by CMP, and by none of the other nucleotide-monophosphates ^29^. CMP binding affinities (*K*_d_) of CST_ZM_ and *human* CST (hCST) were further found to be similar to one another and consistent with Isothermal Titration Calorimetry measurements ^29^. Indeed, we were able to confirm robust transport of CMP-sialic acid in CST_ZM_ containing proteoliposomes comparable to hCST (Fig. 1b). Thus, we could establish CST_ZM_ as a suitable model system to elucidate the molecular basis of human CMP-sialic acid transport.

CST_ZM_ was crystallized by vapor diffusion and *in meso* (LCP) methods in the absence and presence of CMP, respectively (Methods). The CST_ZM_ crystal structures were solved by molecular replacement (MR) and refined against data extending up to 3.4 Å and 2.8 Å, respectively (Supplementary Fig. 1b, Supplementary Table 1, and Methods). Both CST_ZM_ structures were captured in an outward-facing conformation, which in a cellular context places the cavity open to the Golgi lumen for the binding of CMP (Fig. 1a and 1c). As expected from the low sequence identity, a comparison of CST_ZM_ with the previous outward-facing structure of the GDP-mannose *yeast* transporter Vrg4 showed a large degree of structural difference, with an r.m.s.d. of 4.9 Å for 179 pairs of Cα atoms (Methods and Supplementary Fig. 2a). The poor superimposition was also observed in the *apo* CST_ZM_ and Vrg4 structures, which means that the structural differences were unlikely to be the consequence of binding different substrates (Methods and Supplementary Fig. 2b). Rather, extensive re-positioning of TM3, TM6 and TM8, in particular, reshapes the outward-facing state to give uniquely different cavities for CST_ZM_ and Vrg4 (Fig. 1e and Supplementary Fig. 2a).

Due to the destabilization of CST_ZM_ by monoolein, LCP crystals could only be obtained in the presence of 1.2 mM CMP, which significantly stabilized the protein ^29^. Consistently, an omit map showed strong positive density for CMP, which was absent in the *apo* CST_ZM_ structure obtained by vapor diffusion crystals (Supplementary Fig. 2a). Unexpectedly, CMP binds at the peripheral end of the large cavity, between the helices TM2, TM3 and TM7 (Fig. 1c-e). The binding site for CMP shares no overlap with the position of GDP in the GDP-mannose bound structure of Vrg4, where the substrate binds more centrally and is coordinated by residues from TM1, TM7, TM8 and TM9 (Fig. 1e and Supplementary Fig. 2). Rather, the repositioning of TMs in CST_ZM_ creates a unique pocket for CMP not observed in Vrg4 (Fig. 1e and Supplementary Fig. 2a). Reflecting their poor overall structural superimposition, the side-chain chemistry between the two proteins is highly divergent at the respective GDP-mannose and CMP binding sites (Supplementary Fig. 3a-b). Despite these differences, most residues surrounding CMP within the CST subfamily are highly conserved (Supplementary Fig. 3c).

### Molecular recognition of CMP and CMP-sialic acid by CST_ZM_ and human CST

Using the GFP-TS assay, we found that the addition of CMP increased the melting temperature (*T*_*m*_) of CST_ZM_ by ~7 °C compared to only ~ 1° C for either UMP or sialic acid, and was hence used to measure substrate binding (Supplementary Fig. 1c-d). In the CST_ZM_ structure, the nucleobase cytosine forms π-stacking interactions with Trp209 and hydrogen bonds to the carbonyl group of Asn86 (Fig. 2a). The ribose moiety is coordinated by hydrogen bonds to Ser79 and by van der Waals interactions to Leu83 and Phe205. The phosphate group is coordinated to the side chains of Glu42 and Try82 *via* highly ordered water molecules. With the exception of Ser79, single alanine mutants of these residues strongly reduced CMP binding (Fig. 2b). The alanine mutants that completely abolished CMP binding were Tyr82, Asn86, and Trp209. Apart from Trp209, all these residues are strictly conserved in related SLC35 members, but are not conserved in Vrg4 at either the sequence or structural level (Supplementary Fig. 3a-b and Supplementary Fig. 4a-b). In support of their relevance to CST proteins, single alanine substitutions at equivalent positions in hCST also abolished CMP binding (Fig. 2b). As in CST_ZM_, the equivalent serine to alanine mutation in hCST (Ser95) showed robust binding. Thus, polar interactions with ribose do not appear critical for CMP binding, rather CMP binding appears to be primarily established through coordination of the phosphate and cytosine moieties.

**Fig. 2.**
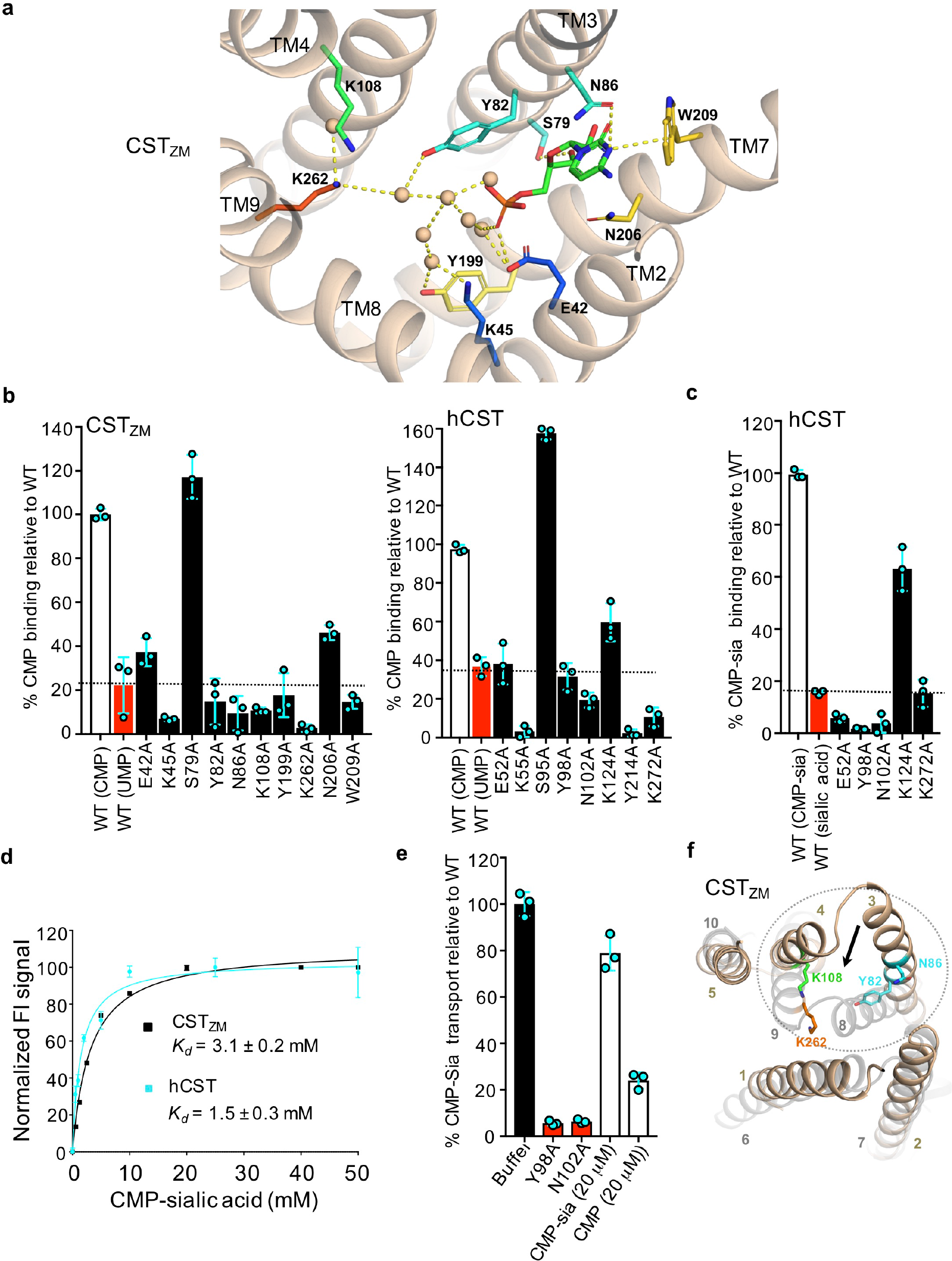
Molecular recognition of CMP by CST_ZM_. **a**, The substrate-binding site in the outward facing CST_ZM_ structure in complex with CMP (TMs in light brown). The CMP is shown as green sticks, waters as light-brown spheres, and residues coordinating CMP and waters are shown as sticks (colored as in Fig. **1d**). Dashed lines represent hydrogen-bonds distances at ~2.7 to 3.5Å, except the longer K262 to water distance of 4.3Å; water positions are conserved between the two molecules in the a.s.u. within 1.7 Å. The π-cation interaction between W209 and cytosine nucleobase is also shown. Single alanine mutants of hCST residues equivalent to Y199 (Y214^57,58^), and K262 (K272^59^) have previously shown to be essential for CST activity. **b**, CMP binding to CST_ZM_ (left panel) and hCST (right panel) using CST-GFP-fusions as determined by a shift in Δ*T*_*m*_ after addition of 1 mM CMP to WT (open bars) and to mutants (filled bars). Non-specific binding was estimated with addition of 1 mM UMP (red bars). **c**, CMP-sialic acid binding to hCST using hCST-GFP-fusions as determined by a shift in Δ*T*_*m*_ after addition of 10 mM CMP-sialic acid to WT (open bars) and to mutants (filled bars). Non-specific binding was estimated after addition of sialic acid (red bars). **d**, The GFP-TS assay was used to estimate the binding affinity of CMP-sialic acid to CST_ZM_ (black squares) and hCST (cyan squares). Binding affinities (*K*_d_) were calculated using data from a range of CMP-sialic acid concentrations, and were fitted by non-linear regression using data from 3 independent experiments (the values reported are the mean ± s.e.m. of the fit; see Methods). **e**, Competitive uptake of [^3^H]-CMP-sialic acid by hCST in proteoliposomes pre-incubated with CMP in the presence of either buffer (filled) or 20 μM CMP/CMP-sialic acid (non-filled). Non-specific transport was estimated from the data of the non-CMP-sialic acid binding mutants Y98A and N102A shown in C. (red). In all experiments errors bars, s.e.m.; n = 3. **f**, Superimposition of the structural-inverted repeats TMs 1 to 5 (cartoon; light brown) and TMs 6 to 10 (cartoon; light grey); asymmetry of structural symmetrical elements identifies local, gating elements ^23, 60, 61^. Side-chains of Lys108, Ty82 and Asn86 involved in CMP-sialic acid binding are shown as sticks (colored as in Fig. **1d**).

In a surface representation of CST_ZM_, the phosphate moiety points towards a large patch of positively-charged electrostatic potential, which is a highly favorable interaction surface for the negatively charged sialic acid sugar (Fig. 1c). In addition to the waters interacting directly with the phosphate moiety, there is also a network of well-coordinated waters that extends from one end of the 23 Å cavity to the other (Fig. 2a and Supplementary Fig. 3d). Noticeably, crystallographic waters at the opposite end of the cavity to CMP are connected to well-conserved lysine residues, Lys108 and Lys262 (Fig. 2a and Supplementary Fig. 4). Remarkably, lysine to alanine substitutions abolished CMP binding in CST_ZM_ and in hCST the equivalent residue to Lys262 (Lys272) was abolished whilst Lys108 (Lys124) showed reduced binding (Fig. 2b). In comparison, alanine substitutions of the polar residues Asn106 and Ser267 located near these residues, but not interacting with these waters, retained robust CMP binding (Supplementary Fig. 1e). CMP is further connected to the bottom of the cavity through water-mediated contacts to Lys45 and Tyr199, which are conserved in hCST (Lys55, Tyr214) and were likewise found to be essential for binding (Fig. 2b). Thus, the binding of CMP is additionally coordinated by an intricate hydrogen-bonding network of waters and charged groups.

One would expect that the binding site for CMP and for CMP when it is conjugated with sialic acid to be equivalent. To confirm this assumption, single alanine mutants assessed for CMP binding were re-measured with CMP-sialic acid. Consistently, CMP and CMP-sialic acid binding profiles in both hCST and CST_ZM_ proteins were almost identical (Fig. 2c and Supplementary Fig. 1d). This finding strongly suggests that the highly-coordinated waters next to the phosphate moiety likely represent the binding site of the sialic acid moiety when it is conjugated to CMP (Fig. 2a). Modelled in this position, the lysine residue Lys45 would be suitably located to interact directly with sialic acid, whereas Lys108 and Lys262 would indirectly coordinate the sialic acid moiety, but not the phosphate moiety itself (Supplementary Fig. 5a). Notably, there is no obvious overlap in the sugar binding sites for sialic acid in CST and mannose in the GDP-mannose bound structure of Vrg4 (Supplementary Fig. 2a).

### Dimerization and structural impact of human CST disease mutations

An overlay of the internal structural repeats in CST_ZM_ shows that Lys108 and Lys262, along with the CMP binding residues Asn86 and Tyr82, are consistently located on the segments most likely to undergo local rearrangements in response to substrate binding (Fig. 2f) ^30^. Furthermore, CST_ZM_ homodimerizes in both crystal forms through extensive interactions between equivalent TM5 and TM10 helices, burying a total surface area of ~3.060Å^2^ (Fig. 3a). CST_ZM_ appears to be forming two distinct domains, a dimerization domain and a transport domain. The transport domain bundles are held together on the cytoplasmic side through extensive polar interactions (Supplementary Fig. 5b). It is likely that the dimerization domain acts as a scaffold to support the transport domain bundles rearrangements, which are likely to be similar to that predicted for Vrg4 ^23^ and structural homologues TPT ^30,31^ and YddG ^32^.

**Fig. 3.**
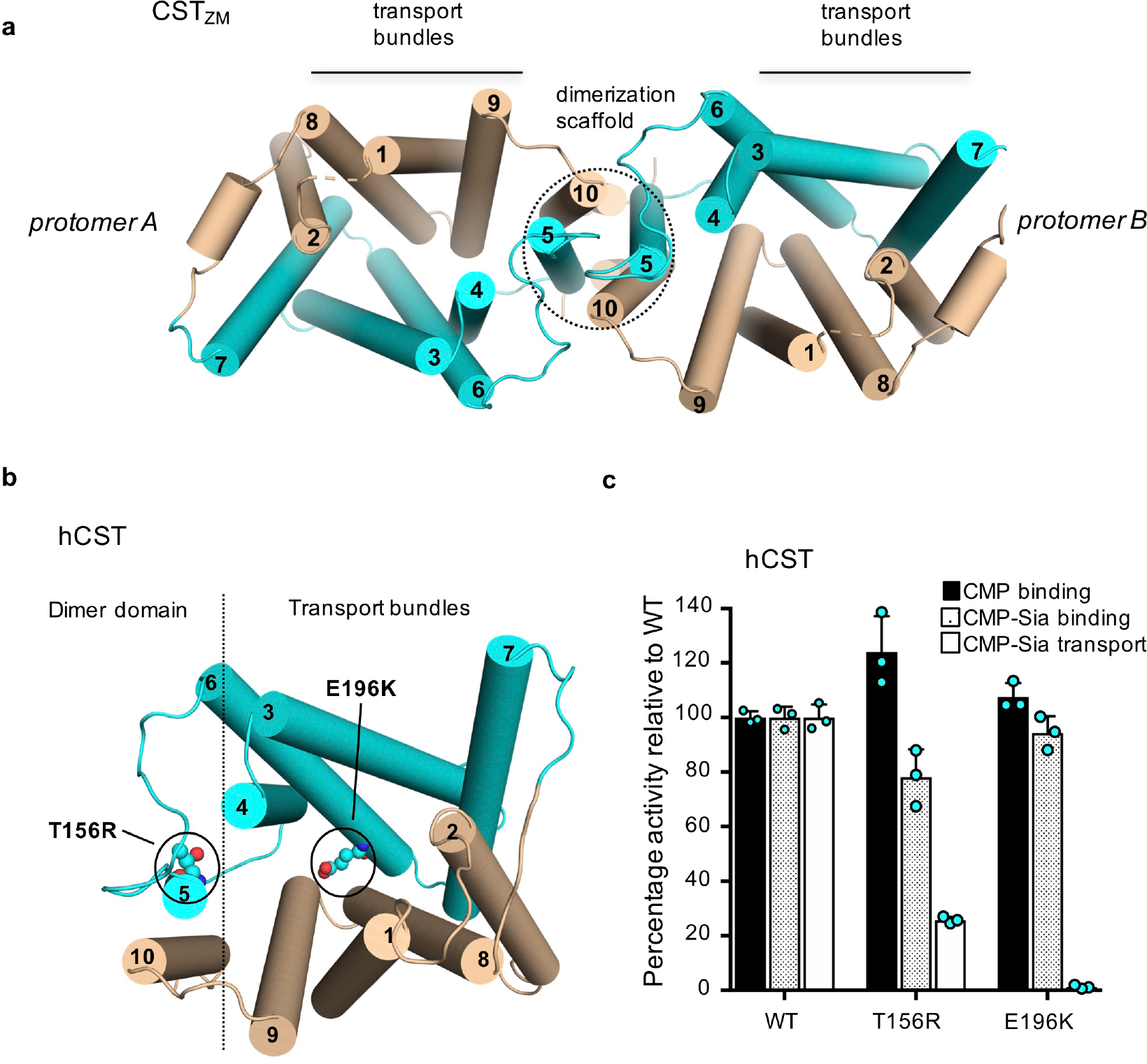
Dimerization and human CST glycosylation disease mutants. **a**, Cartoon representation of the CST_ZM_ homodimer, which is identical in both crystal forms (for sake of clarity only the dimer structure grown from LCP crystals is shown). In each protomer, dimerization is mediated by TM5 (cyan) and TM10 (wheat), which are connected *via* flexible loops to the transport bundles made up of TMs 3, 4, 6, and 7 (cyan) and TMs 1, 2, 8 and 9 (wheat). During structural transition from the outward to occluded conformation the extracellular half of TM 3 and 4 moves inwards (from residue Pro78 to Gly113) to gate substrate binding, as apparent from the asymmetry of these regions in a superimposition of the inverted-topology repeats (Fig. **2f**), which is consistent with what was previously described for other drug-metabolite superfamily (DMT) members ^23,30–32^. In contrast to other dimeric DMT structures ^23,30^, homodimerization of CST_ZM_ is more extensive and not mediated by any lipids. **b**, Cartoon representation of a modelled hCST protein (see Methods) depicting the location of mutations responsible for congenital disorders of glycosylation type 2f. **c**, CMP-sialic acid binding to hCST using hCST-GFP-fusions as determined by a shift in Δ*Tm* after addition of either 1 mM CMP (black bars) or 10 mM CMP-sialic acid (pattern-filled bars) for WT, T156R and E196K mutants. [^3^H]-CMP-sialic acid 2 min uptake by hCST in proteoliposomes pre-incubated with CMP for WT, T156R and E196K (open bars). In all experiments errors bars, s.e.m.; n = 3.

CST are associated with Congenital disorders of glycosylation type 2f (CDG2f), which result in severe inherited diseases caused by under-sialyation of glycoproteins ^18,33,34^. More specifically, heterozygous mutations Thr156Arg and Glu196Lys in CST have been shown to reduce CMP-sialic acid transport into Golgi ^18^. In a model of hCST based on CST_ZM_, Thr156 is located at the dimer interface and Glu196 is located at the cytoplasmic intracellular ionic network (Fig. 3b). Using the GFP-TS assay we confirmed that the Thr156Arg and Glu196Lys mutations show close to wildtype binding of CMP and CMP-Sia, which means these mutations have not perturbed the hCST structure (Fig. 3c). Rather, CMP-Sialic acid transport by hCST was severely reduced in the Thr156Arg disease mutation and abolished in Glu196Lys (Fig. 3c). Thus, the disease mutations support the physiological importance of CST dimerization and the intracellular ionic network.

### A chaperone-mediated nucleotide substrate selectivity mechanism

Although the positively charged binding site for sialic acid is highly favorable, it has been reported that “free” sialic acid does not compete for the transport of CMP-sialic acid ^35^. Consistently, we were unable to detect the binding of sialic acid to CST_ZM_ and hCST proteins (Fig. 2c and Supplementary Fig. 1d). We think that the most likely reason is that the sialic acid moiety is unable to reach the deep binding pocket on its own. In agreement with this line of reasoning, the sialic acid binding pocket is situated below a large region of non-polar and negatively charged surfaces, which are highly conserved and conspicuously absent of crystallographic waters (Fig. 1c and Supplementary Fig. 3c). In support of a physical barrier for the passage of free sialic acid, we found that the binding affinities (*K*_d_) for CMP-sialic acid (*K*_d_ = 1.5 to 3.1 mM) were significantly weaker than for CMP (*K*_d_ = 40 μM) (Fig. 2d and Supplementary Fig. 1f). Moreover, pre-steady state concentrations of unlabeled CMP competed significantly better for the uptake of radiolabeled [^3^H]-CMP-sialic acid than equimolar amounts of unlabeled CMP-sialic acid (Fig. 2e).

It has been shown that the closely related UDP-galactose transporter (SLC35A2) (Supplementary Fig. 4a), could be converted into a CMP-sialic acid transporter when their respective TM7 segments were swapped, with further enhancement in activity if TM2 and TM3 were also exchanged ^36^. Consistently, we found that only residues from exactly these TMs coordinated CMP in the CST_ZM_ structure (Fig. 2a). Based on the CST_ZM_ structure, sugar binding residues are therefore unlikely to be altered by these chimera studies, which further implies that sugar recognition does not require strict coordination, but is likely to be flexible. Promiscuity could be important *in vivo* for other NST members too, since different sugars can be transported when coupled to the same nucleotide ^37,38^. Additionally, the physiological substrate for CST_ZM_ is likely to be CMP-Kdo, which is also an acidic sugar different yet structurally related to sialic acid ^26^. Taken together, we conclude that the specificity for CMP-sialic acid in CST is primarily established by its recognition of the CMP moiety.

### Human CST and CST_ZM_ can function both as a uniporter and an antiporter

It is well-established that NST function as antiporters, exchanging the cytosolic nucleotide-sugar for their corresponding nucleotide monophosphate ^8,34^. Most antiporters work by simple competition mechanism between structurally similar, yet distinct solute molecules ^39^. This conceptually poses a problem in the NSTs however, as whilst the bi-partite molecule is matched with a bi-partite substrate-binding site, the counter substrate is only a competitor for one half of the transported solute. This “flaw” is most pronounced in CST, since CMP is transported in both directions with substrate recognition governed primarily by the nucleotide monophosphate. Intriguingly, transport measurements in either isolated Golgi vesicles or proteoliposomes concluded that CST activities might be “leaky”, because CMP-sialic acid uptake could be observed in the absence of vesicles pre-loaded with CMP ^8,17,40^. Although compelling, high-levels of CMP-sialic acid uptake into empty vesicles has made it difficult to come to any firm conclusions regarding coupling. To investigate this observation more thoroughly, we used our proteoliposome transport assay that was extensively optimized to have very low background activity in either empty or inactive-mutant containing liposomes (Fig. 1b and Fig. 2e). As shown in Figure 4a, robust transport of [^3^H]-CMP-sialic acid was indeed measurable in the absence of CMP. Transport of [^3^H]-CMP-sialic acid was, however, highly stimulated when liposomes were preloaded with CMP, consistent with other studies and antiport activity (Fig. 4b) ^17^. Remarkably, we found that transport of [^3^H]-CMP (uptake) was also measurable in the absence of an outwardly-directed CMP-sialic acid gradient, consistent with a facilitator primarily recognizing the CMP moiety (Supplementary Fig. 5c-d).

**Fig. 4.**
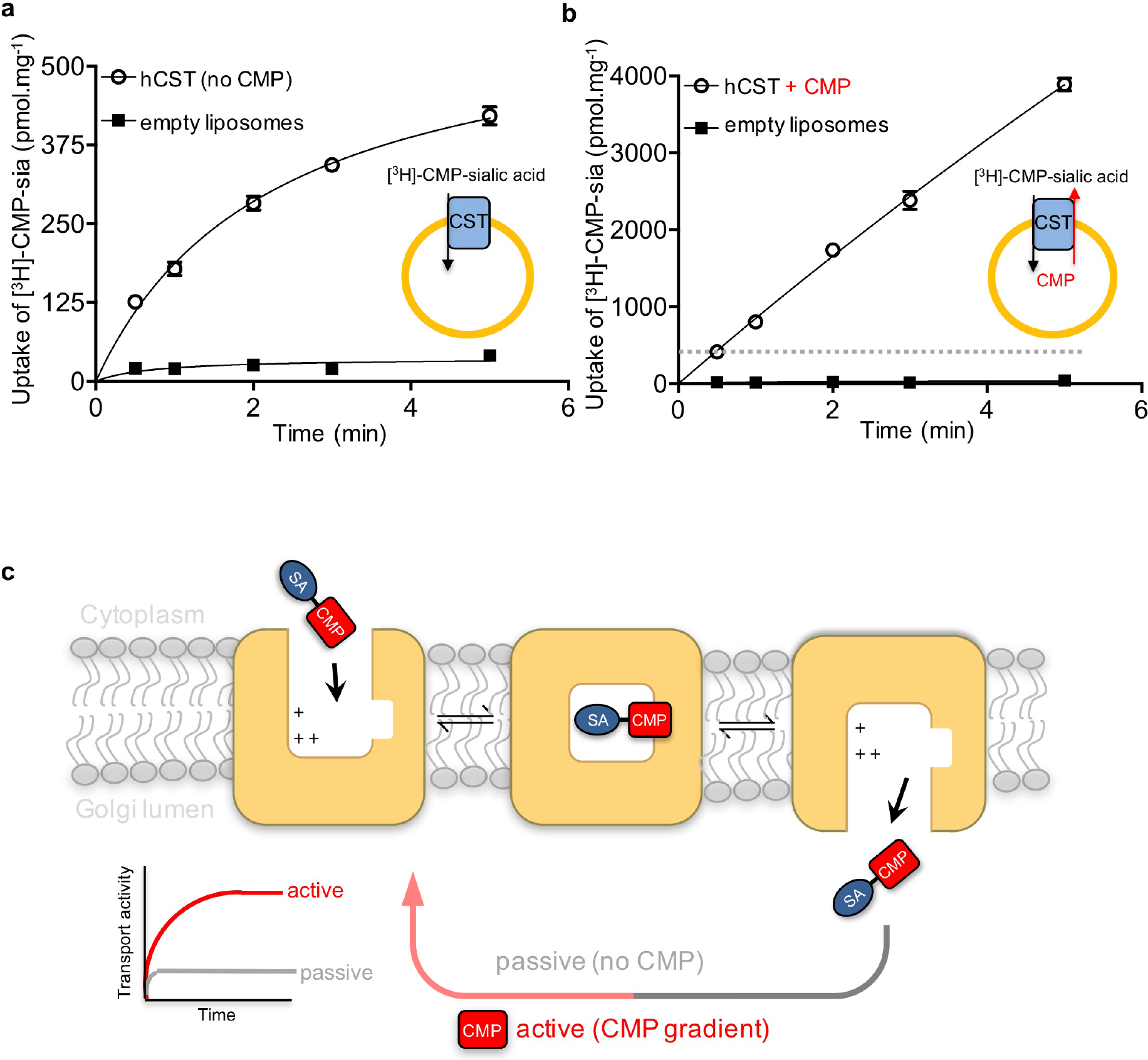
CST operates both as a passive and as an active transporter. **a**, Time-dependent uptake of[^3^H]-CMP-sialic acid by hCST (non-filled circles) in proteoliposomes containing no CMP. The proteoliposome setup is shown as a schematic in the bottom-right hand corner. Non-specific uptake was estimated from data of liposomes containing no protein (filled squares). **b**, As in Figure **4a**. except liposomes were preincubated with CMP. The dashed line represents the maximum total amount of CMP-sialic acid uptake recorded in the absence of CMP, shown in **4a**. Errors bars represent s.e.m. of 3 independent experiments. **c**, CST transporters operate by a rocking-switch alternating-access mechanism as they exchange CMP-sialic acid for CMP across the Golgi membrane using an antiport mechanism ^39^. However, as the recognition for CMP-sialic acid is established primarily by CMP, they can also carry out factitive transport of CMP, which means they can operate both as passive and as active transporters.

## Discussion

By a combination of crystal structures, binding and transport assays we conclude that the molecular recognition of CMP-sialic acid in CST is established primarily through the nucleotide monophosphate CMP. We further propose that sialic acid needs to be chaperoned into the deep binding site by CMP, which requires passage through a cavity that is very different to that seen in the GDP-mannose transporter Vrg4. Indeed, the Nucleotide Sugar Transporters appear to be adopting unique cavities to match the binding modes of the different substrates. While this is a common practice in GPCRs ^41^, this level of divergence is unusual in transporters, which predominantly evolve only the side-chain chemistry to bind different substrates ^39^. This specialization contributes to a nucleotide-mediated selectivity mechanism in CST, which might be to ensure that CMP-sialic acid is not inadvertently transported into the ER by the other nucleotide-sugar transporters, and would explain why CMP is specifically conjugated to sialic acid.

Nucleotide-sugar transporters are annotated as antiporters ^8, 10, 23, 42^, but uniquely we conclude that CMP-sialic acid transporters are antiporters that can also function as facilitative transporters (uniporters) (Fig. 4c). After substrate release, uniporters are able to spontaneously re-set themselves back to an opposite-facing conformation, whilst antiporters are unable to reset themselves until a counter-substrate has bound ^39^. This distinction implies that the energetic barriers separating the alternating conformations in CMP-sialic acid transporters are lower than in other NSTs, which are mostly strict antiporters and are thought to bind their substrates with equal affinities ^23^. From a physiological perspective, CMP efflux might be required to be uncoupled from CMP-sialic acid transport under some circumstances because low levels of CMP (K_i_ = 40 μM) is a potent inhibitor of siayltransferase activity in the Golgi ^43,44^. Indeed, for some unknown reason CMP-sialic acid is synthesized in the nucleus while all other nucleotide-sugars are synthesized in the cytoplasm ^8,45^. The compartmentalization of CMP-sialic acid synthesis in the nucleus suggests that cytoplasmic levels of CMP-sialic acid are regulated, which could be further correlated with the different modes of transport seen in CST. Further investigation will clearly be required to solidify the physiological significance of the uncoupled CMP-sialic acid transport and to refine the conserved sugar and nucleotide recognition motifs for this diverse SLC35 transporter family. Nonetheless, the structure of CST_ZM_ and comparative analysis with *human* CST provides a template for glycopathway engineering, a molecular explanation for their substrate recognition, and explains how they are uniquely able to transport in an energized and non-energized manner.

## Acknowledgments

We are grateful to Yurie Chatzikyriakidou for generation of Figure 1a, and Magnus Claesson for assistance with data collection and manuscript preparation. We thank The European Synchrotron Radiation Facility and Diamond Light Source synchrotrons and the excellent assistance of the beamline scientists. This work was funded by the Knut and Alice Wallenberg Foundation (D.D). D.D acknowledges support from the Wenner-Gren foundation and EMBO through the Young Investigator Program (YIP).

## Author Contributions

D.D. designed the project. Purification and crystallization of CST_ZM_ was carried out by E.N. Data collection was carried out by E.N., M.C., and D.D. Structure determination and refinement of CST_ZM_ was carried out by A.G. Binding experiments were carried out by E.N., and A.G. Transport assays were carried out by A.Q. The manuscript was prepared by D.D and A.G. with assistance from E.N, A.Q. and M.C. All authors discussed the results and commented on the manuscript.

## Competing interests

The authors declare no competing interests.

## Supplementary Information

Supplemental Information includes five figures and one table

## Methods

### Target identification using fluorescence-based screening methods

CST homologues were synthesized as gene-strings (Life Technologies) and were cloned into the GAL1 inducible TEV cleavable GFP-His_8_ 2μ vector **pDDGFP2**, and transformed into the *S. cerevisiae* strain FGY217 (MATα, ura3–52, lys2Δ201, and pep4Δ)^46^ and overexpressed as previously described ^27,47^. In brief, 10 ml *S. cerevisae* FGY217 cell cultures in -URA media and 0.1% (v/v) glucose were grown at 30°C and expression was induced by the addition of final 2% (v/v) D-galactose at an OD_600_ of 0.6. After ~22 hrs, cells were harvested, resuspended in 1 × PBS buffer and overexpression levels assessed by whole-cell GFP fluorescence ^27,47^. Fusions with detectable expression levels were re-grown in larger 2L culture volumes and membranes subsequently isolated. The monodispersity of expressed fusions were screened by fluorescence-detection size exclusion chromatography (FSEC)^28^ as described previously ^47^. Out of the CST homologues screened, CST from *Zea mays* (CST_ZM_: UniProt accession number: B4FZ94) was the most stable after purification in detergent ^48^. To facilitate crystallization of CST_ZM_, five non-conserved loop residues were mutated using overlap PCR based on a close homologue from *Fragaria vesca*, which was predicted to be less disordered: His58Arg, Ser59Thr, Ser61Pro, Pro62Ser and Pro63Val. C-terminal residues retained after TEV cleavage are shown in italic. Unless otherwise stated this wildtype-like construct was used for all experiments.

MQWYLVAALLTILTSSQGILTTLSQSNGKYNYDYATIPFLAELFKLSVSGFFLWKEC<u>RT</u>S <u>PSV</u>RMTKEWRSVRLYVVPSVIYLIHNNVQFATLTYVDPSTYQIMGNLKIVTTGILFRLVL KRKLSNIQWMAIVLLAVGTTTSQVKGCGDSPCDSLFSAPLEGYLLGILSACLSALAGVYTEYLMKKNNDSLYWQNVQLYTFGVIFNMGWLIYGDFKAGFELGPWWQRLFNGYSITTWMVVFNLGSTGLLVSWLMKYSDNIVKVYSTSMAMLLTMVLSIYLFSVKATIQLFLGIIIC IISLQMYFMPVHMLIELPQTLPVTSK*ENLYFQ*

### Large-scale production and purification of CST_ZM_ and hCST

Cells were harvested from 12 L *S. cerevisiae* cultures, resuspended in buffer containing 50 mM Tris-HCl pH 7.6, 1 mM EDTA, 0.6 M sorbitol, and lysed by mechanical disruption as described previously ^27^. Membranes were isolated by ultracentrifugation at 195,000 *g* for 2 h, homogenized in 20 mM Tris-HCl pH 7.5, 0.3 M sucrose, 0.1 mM CaCl2, flash-frozen in liquid nitrogen and stored at −80 °C. Membranes were resuspended in 200 ml of equilibration buffer (EB) consisting of 1 × PBS, 150 mM NaCl, 10 % (v/v) glycerol. Membranes were solubilized with 2 % (w/v) DDM for 1h at 4 °C. Protein was purified in DDM as described previously ^27^. In brief, after ultracentrifugation at 195,000 *g* for 45 min, the supernatant (~200 ml) was supplemented with 10 mM imidazole pH 7.5 and incubated with 10 mL of Ni^2+^-NTA resin for 2h at 4 °C. The resin was washed with 200 mL of EB supplemented with 30 mM imidazole pH 7.5 and 0.1 % (w/v) DDM, and the protein was eluted using 20 mL of EB containing 250 mM imidazole pH 7.5 and 0.1 % (w/v) DDM. Equimolar TEV-His6 protease was added to the elution and dialyzed o/n in 3L of buffer containing 10 mM Tris-HCl, pH 7.5, 150 mM NaCl, and 0.03 % (w/v) DDM. After proteolytic removal of GFP-His_8_ by TEV-His_6_ protease the material was loaded onto a 5 mL HisTrap column (no imidazole), and the flow-through collected. The untagged CST_ZM_/hCST proteins were further purified by SEC in buffer containing 20 mM Tris-HCl, pH 7.5, 150 mM NaCl, and 0.03 % (w/v) DDM, and then concentrated to ~3 mg.ml^−1^ for functional experiments.

### Transport activity of reconstituted hCST/CST_ZM_

Total bovine brain lipid extract and cholesteryl-hemisuccinate were mixed in buffer containing 10 mM Tris-HCl pH 7.5 and 2 mM MgSO_4_ to reach a final concentration of 30 and 6 mg.mL^−1^, respectively. Either non-deuterated CMP at 0.5 mM or non-deuterated CMP-sialic acid at a final concentration of 1.5 mM were added prior to forming liposomes by multiple rounds of freeze-thaw cycles by flash-freezing in liquid nitrogen, and thawing at room temperature and sonication. To make proteoliposomes, 10 μg of purified hCST/CST_ZM_ was added into 500 μL of unilamellar vesicles, flash frozen and then thawed at room temperature. Large, unilamellar proteoliposomes were prepared by extrusion through a filter with membrane pore size of 400 nm. For removal of excess cold substrate, proteoliposomes were pelleted by ultracentrifugation at 150000 *g* for 1 hr, the supernatant removed, and the pellet resuspended in Tris-MgSO_4_ pH 7.5 buffer to a final lipid concentration of 80 mg.mL^−1^. For transport measurements, each reaction contained 5 μL of proteoliposomes that were diluted into 45 μL of Tris-MgSO_4_ pH 7.5 buffer containing either radiolabeled 0.05 μM [^3^H]-CMP or 0.2 μM [^3^H]-CMP-sialic acid (American Radiolabeled Chemicals, Inc). Reaction was stopped by addition of 1 ml of Tris-MgSO_4_ buffer, filtered, and further washed with 6 mL of the same buffer. Non-specific uptake was estimated from empty or non-substrate-binding mutant containing liposomes. The radioactivity corresponding to the internalized substrate was measured by scintillation counting. Each experiment was performed in triplicate.

### Substrate specificity and binding analysis using GFP-TS

The GFP-TS method, which is derived from the FSEC-TS assay ^49^, was used to screen for ligand binding as described previously ^29^. In brief, membranes harboring either hCST or CST_ZM_ (wildtype sequence) were diluted to a final concentration of 3.5 mg.ml^−1^ in 30 mL of buffer containing 150 mM NaCl, 20 mM Tris-HCL pH 7.5, and 1 % DDM. After 1-hour solubilization in DDM at 4°C the detergent-solubilized membranes were divided into 4 mL aliquots. The compounds CMP, UMP, TMP and GMP dissolved in pure water were separately added to each of the 4 mL aliqouts to reach a final concentration of 1 mM. The compounds sialic acid (pH adjusted to 7.5) and CMP-sialic were freshly prepared in pure water and added to the remaining aliquots to reach a final concentration of 10 mM. After addition of final 1 % (w/v) β-octyl-D-glucoside (OG), 120 μL fractions were aliquoted into PCR tubes. The PCR tubes were heated at individual temperatures ranging from 4 to 100 °C for 10 minutes, and aggregated material was sedimented by centrifugation at 5,000 *g* for 30 mins at 4 °C using a Microfuge^®^ 18 Centrifuge, Beckman Coulter. The resulting supernatants were transferred into a 96-well black clear bottom plate (Nunc) and the GFP fluorescence (excitation/emission wavelengths: 488 nm/512 nm) intensity measured using a SpectraMax Germini EM microplate reader (Molecular Devices). For control samples pure water was added instead of compounds and were treated in the same way. Each experiment was carried out in triplicate. The hCST/CST_ZM_ melting curves (*T_m_*) were fitted by non-linear regression using the software Prism. The Δ*T*_*m*_ was calculated by subtracting the apparent *T*_*m*_ in the presence and absence (control) of the aforementioned compounds.

For binding affinity measurements by GFP-TS, solubilized membrane hCST/CST_ZM_ fusions were prepared as described for the *Substrate specificity and binding analysis*, and investigated at 10 different final concentrations of CMP-sialic acid ranging from 0 to 50 mM. The prepared protein samples were incubated for 10 min at 5 °C above their respective apparent melting temperatures, and subjected to centrifugation at 5,000 *g* for 30 mins at 4 °C using a Microfuge^®^ 18 Centrifuge, (Beckman Coulter). The *K_d_* values were calculated from the data points of the 10 different CMP-sialic concentrations, fitted by nonlinear regression (one site, total binding), and the values reported are the averaged mean ± s.e.m. of the fit from n = 3 independent titrations.

CMP binding affinities to detergent-solubilized hCST/CST_ZM_ GFP fusion containing membranes were previously assessed using the GFP-TS assay ^29^. To verify that these CMP binding affinities were comparable to those measured with purified hCST/CST_ZM_ GFP fusions, the experiment was performed as described for CMP-sialic acid, except CMP was added to final concentrations of 0 to 2 mM and the protein concentration was adjusted to 5,000 RFU (0.04 μg.μl^−1^) in 20 mM Tris-HCL pH 7.5, 150 mM NaCl, and 0.03 % (w/v) DDM.

### Analysis of the substrate-binding site

All CST mutants were synthesized as gene strings (Life Technologies) and were sub-cloned, overexpressed and membranes were isolated as described in the section *Large-scale production and purification of CST_ZM_ and hCST*. Only hCST/CST_ZM_ mutants with a monodisperse peak in DDM by FSEC ^28^ were assessed for their ability to bind CMP or CMP-sialic acid using the GFP-TS assay. The apparent *T*_*m*_ was determined in the absence and presence of compounds (CMP, UMP, CMP-sialic acid and sialic acid), as described for the substrate-specificity measurements. All melting curves were fitted by non-linear regression using the software Prism. Each experiment was carried out in triplicate. The Δ*T*_*m*_ was calculated by subtracting the absolute difference in the melting temperatures (*T*_*m*_) in the presence and absence of listed compounds. Because the binding affinities measured by the GFP-TS method were equivalent to binding affinities estimated by ITC ^29^, the Δ*T*_*m*_ values were considered a robust estimate of substrate-binding efficiency. The comparative binding efficiency of mutants was calculated as fraction of WT Δ*T*_*m*_hCST/CST_ZM_ values. Y98A and N102A mutants of hCST were purified as described for the wildtype and concentrated to ~3 mg.ml^−1^ for functional experiments.

## Crystallization of CST_ZM_

The initial purification steps, up until binding of DDM solubilized CST_ZM_ fusion containing membranes, was performed as described in the section *Large-scale production and purification of CST_ZM_ and hCST*. The binding of DDM solubilized membranes was carried out by incubation with 5 mL of Strep-Tactin^®^XT resin (Cat. 2-4010-025; IBA GmbH, Germany) for 2h at 4 °C. The collected resin was transferred into a 100 mL Econo column and washed with 100 mL of buffer containing 1 × PBS, 150 mM NaCl, 10 % (v/v) glycerol, and 0.45 % (w/v) nonyl-β-D-maltopyranoside (NM). The Strep-Tactin resin containing immobilized CST_ZM_-GFP-His_8_-Strep_tag_ fusion protein was transferred into a 50 mL falcon tube, 5 mg of recombinant His_6_-tagged TEV protease was added and incubated o/n at 4°C under gentle shaking for proteolytic digestion. The affinity resin was sedimented by centrifugation and the resulting supernatant containing the untagged CST_ZM_ protein and His-tagged TEV protease was passed through a 5 mL HisTrap column (GE Healthcare) pre-equilibrated with buffer composed of 20 mM Tris-HCl pH 7.5, 150 mM NaCl, 0.45 % (w/v) NM. The flow-through was collected and concentrated (100 MWCO). The protein was subjected to size-exclusion chromatography (SEC) using a Superdex 200 column (GE Healthcare) in buffer containing 20 mM Tris-HCl pH 7.5, 150 mM NaCl, 0.45 % (w/v) NM. Initial vapour-diffusion crystals of purified CST_ZM_ at ~5 to 10 mg.mL^−1^ (0.2, 0.2 nl drop size) was obtained using the MemGold^TM^ screen (Molecular Dimensions). Following crystallization optimization, improved crystals were obtained by the hanging drop method against 500 μL of reservoir solution containing 35 % (v/v) PEG 400, 0.1 M NaCl, 0.1 M MES (pH 6.5). Crystals were grown in 2 μL drops containing 10 mg.mL^−1^ purified CST_ZM_ and reservoir solution in 1:1 ratio. Crystals appeared after 1 day and reached maximum size in 4 ~days. For data collection crystals were flash-frozen and stored in liquid nitrogen.

To obtain a substrate bound structure, CMP was added to the CST_ZM_ protein solution to a final CMP concentration of 1.2 mM. LCP crystals were grown by mixing CST_ZM_ (20 to 40 mg/mL) with liquid monoolein (Sigma, CAS No.111-03-5) in a weight ratio of 2:3 respectively, using coupled syringe-mixing device (Hamilton). Crystallization was achieved by dispensing 50 nL cubic phase bolus onto a 96-well Laminex glass plate (MD11-50, Molecular Dimensions). The bolus was covered with 800 nL of precipitant solution containing 0.1 M sodium chloride, 0.4 M ammonium sulfate, 0.1 M lithium sulfate, 0.1 M sodium citrate tribasic dehydrate pH 5.0, 3 % (v/v) tert-butanol and 30 % (v/v) PEG 300, using an LCP Mosquito crystallization robot (TTP Labtech). Plates were sealed with a Laminex glass cover (MD11–52, Molecular Dimensions) and stored at 20 °C. Crystals appeared after one day and grew to maximum size in 1 week after which they were harvested and flash-frozen and stored in liquid nitrogen.

## Data collection, structure determination and analysis

X-ray crystallographic diffraction data was collected at the European Synchrotron Radiation Facility (Grenoble, France) and Diamond Light Source (Didcot, Great Britain), and processed using XDS ^50^ and the CCP4 suite ^51^. The data of the CST_ZM_ in complex with CMP belonged to the space group C2, with two protomers per asymmetric unit. Phasing with Vrg4 was unsuccessful. Initial phases were obtained by molecular replacement with Phaser ^52^ using a homology model derived from the bacterial transporter Yddg (PDB id: 5i20), which shares 23 % sequence identity to CST_ZM_. DEN refinement was used for initial phase improvement ^53^ and automated model building was performed using Phenix AutoBuild ^54^. The model of CST_ZM_ in complex with CMP was built by several rounds of refinement using REFMAC5 ^55^ interspersed with model building using COOT ^56^ to *R* and *R*_free_ values of 23.9 and 25.3 %, respectively. For the C-terminal residues 314-322, and the loop residues 27-30 no interpretable electron density was observed, hence these were not modelled. The data from crystals yielding the *apo* structure of CST_ZM_ belonged to the P1 space group, and the asymmetric unit was composed of the physiological dimer. Identical dimers were also observed in LCP crystals after applying a crystallographic two-fold. The *apo* structure was refined to the final R and R_free_ of 26.8% and 28.6%, respectively. Structural superimpositions were performed using the align command of PYMOL software using C-alpha coordinates. Homology model of hCST was generated using Swiss-model server (https://swissmodel.expasy.org) with default settings. CST_ZM_ structure was used as a template. Figures of structures were prepared using PyMOL (http://www.pymol.org/).

## Data availability

The coordinates and the structure factors for CST_ZM_ have been deposited in the Protein Data Bank with accession numbers 6I1R (CMP bound CST_ZM_ structure) and 6I1Z (*apo* CST_ZM_ structure). Further information and requests for resources and reagents should be directed to David Drew.

## References

1. Varki, A. Sialic acids in human health and disease. Trends Mol Med 14, 351–60 (2008).

2. Varki, A. & Schauer, R. Sialic Acids. in Essentials of Glycobiology (eds. nd et al.) (Cold Spring Harbor (NY), 2009).

3. Yoo, S.W. et al. Sialylation regulates brain structure and function. FASEB J 29, 3040–53 (2015).

4. Han, J. et al. Genome-wide CRISPR/Cas9 Screen Identifies Host Factors Essential for Influenza Virus Replication. Cell Rep 23, 596–607 (2018).

5. Bull, C., Stoel, M.A., den Brok, M.H. & Adema, G.J. Sialic acids sweeten a tumor’s life. Cancer Res 74, 3199–204 (2014).

6. Rodrigues, E. & Macauley, M.S. Hypersialylation in Cancer: Modulation of Inflammation and Therapeutic Opportunities. Cancers (Basel) 10 (2018).

7. Bauer, J. & Osborn, H.M. Sialic acids in biological and therapeutic processes: opportunities and challenges. Future Med Chem 7, 2285–99 (2018).

8. Hadley, B. et al. Structure and function of nucleotide sugar transporters: Current progress. Comput Struct Biotechnol J 10, 23–32 (2015).

9. Eckhardt, M., Muhlenhoff, M., Bethe, A. & Gerardy-Schahn, R. Expression cloning of the Golgi CMP-sialic acid transporter. Proc Natl Acad Sci U S A 93, 7572–6 (1996).

10. Hirschberg, C.B., Robbins, P.W. & Abeijon, C. Transporters of nucleotide sugars, ATP, and nucleotide sulfate in the endoplasmic reticulum and Golgi apparatus. Annu Rev Biochem 67, 49–69 (1998).

11. Wex, B. et al. SLC35B4, an Inhibitor of Gluconeogenesis, Responds to Glucose Stimulation and Downregulates Hsp60 among Other Proteins in HepG2 Liver Cell Lines. Molecules 23(2018).

12. Yazbek, S.N. et al. Deep congenic analysis identifies many strong, context-dependent QTLs, one of which, Slc35b4, regulates obesity and glucose homeostasis. Genome Res 21, 1065–73 (2011).

13. Winter, G.E. et al. The solute carrier SLC35F2 enables YM155-mediated DNA damage toxicity. Nat Chem Biol 10, 768–773 (2014).

14. Rowland, A., Mackenzie, P.I. & Miners, J.O. Transporter-mediated uptake of UDP-glucuronic acid by human liver microsomes: assay conditions, kinetics, and inhibition. Drug Metab Dispos 43, 147–53 (2015).

15. Kutsuno, Y., Itoh, T., Tukey, R.H. & Fujiwara, R. Glucuronidation of drugs and drug-induced toxicity in humanized UDP-glucuronosyltransferase 1 mice. Drug Metab Dispos 42, 1146–52 (2014).

16. Caffaro, C.E. et al. A single Caenorhabditis elegans Golgi apparatus-type transporter of UDP-glucose, UDP-galactose, UDP-N-acetylglucosamine, and UDP-N-acetylgalactosamine. Biochemistry 47, 4337–44 (2008).

17. Berninsone, P., Eckhardt, M., Gerardy-Schahn, R. & Hirschberg, C.B. Functional expression of the murine Golgi CMP-sialic acid transporter in saccharomyces cerevisiae. J Biol Chem 272, 12616–9 (1997).

18. Ng, B.G. et al. Encephalopathy caused by novel mutations in the CMP-sialic acid transporter, SLC35A1. Am J Med Genet A 173, 2906–2911 (2017).

19. Martinez-Duncker, I. et al. Genetic complementation reveals a novel human congenital disorder of glycosylation of type II, due to inactivation of the Golgi CMP-sialic acid transporter. Blood 105, 2671–6 (2005).

20. Pagan, J.D., Kitaoka, M. & Anthony, R.M. Engineered Sialylation of Pathogenic Antibodies In Vivo Attenuates Autoimmune Disease. Cell 172, 564–577e13 (2018).

21. Ghaderi, D., Taylor, R.E., Padler-Karavani, V., Diaz, S. & Varki, A. Implications of the presence of N-glycolylneuraminic acid in recombinant therapeutic glycoproteins. Nat Biotechnol 28, 863–7 (2010).

22. Kwak, C.Y. et al. Enhancing the sialylation of recombinant EPO produced in CHO cells via the inhibition of glycosphingolipid biosynthesis. Sci Rep 7, 13059 (2017).

23. Parker, J.L. & Newstead, S. Structural basis of nucleotide sugar transport across the Golgi membrane. Nature 551, 521–524 (2017).

24. Perez, M. & Hirschberg, C.B. Topography of glycosylation reactions in the rough endoplasmic reticulum membrane. J Biol Chem 261, 6822–30 (1986).

25. Martinez-Duncker, I., Mollicone, R., Codogno, P. & Oriol, R. The nucleotide-sugar transporter family: a phylogenetic approach. Biochimie 85, 245–60 (2003).

26. Bakker, H. et al. A CMP-sialic acid transporter cloned from Arabidopsis thaliana. Carbohydr Res 343, 2148–52 (2008).

27. Drew, D. et al. GFP-based optimization scheme for the overexpression and purification of eukaryotic membrane proteins in Saccharomyces cerevisiae. Nat Protoc 3, 784–98 (2008).

28. Kawate, T. & Gouaux, E. Fluorescence-detection size-exclusion chromatography for precrystallization screening of integral membrane proteins. Structure 14, 673–81 (2006).

29. Nji, E., Chatzikyriakidou, Y., Landreh, M. & Drew, D. An engineered thermal-shift screen reveals specific lipid preferences of eukaryotic and prokaryotic membrane proteins. Nat Commun 9, 4253 (2018).

30. Lee, Y. et al. Structure of the triose-phosphate/phosphate translocator reveals the basis of substrate specificity. Nat Plants 3, 825–832 (2017).

31. Takemoto, M., Lee, Y., Ishitani, R. & Nureki, O. Free Energy Landscape for the Entire Transport Cycle of Triose-Phosphate/Phosphate Translocator. Structure 26, 1284–1296e4 (2018).

32. Tsuchiya, H. et al. Structural basis for amino acid export by DMT superfamily transporter YddG. Nature 534, 417–20 (2016).

33. Mohamed, M. et al. Intellectual disability and bleeding diathesis due to deficient CMP--sialic acid transport. Neurology 81, 681–7 (2013).

34. Song, Z. Roles of the nucleotide sugar transporters (SLC35 family) in health and disease. Mol Aspects Med 34, 590–600 (2013).

35. Carey, D.J., Sommers, L.W. & Hirschberg, C.B. CMP-N-acetylneuraminic acid: isolation from and penetration into mouse liver microsomes. Cell 19, 597–605 (1980).

36. Aoki, K., Ishida, N. & Kawakita, M. Substrate recognition by nucleotide sugar transporters: further characterization of substrate recognition regions by analyses of UDP-galactose/CMP-sialic acid transporter chimeras and biochemical analysis of the substrate specificity of parental and chimeric transporters. J Biol Chem 278, 22887–93 (2003).

37. Caffaro, C.E., Hirschberg, C.B. & Berninsone, P.M. Independent and simultaneous translocation of two substrates by a nucleotide sugar transporter. Proc Natl Acad Sci U S A 103, 16176–81 (2006).

38. Berninsone, P., Hwang, H.Y., Zemtseva, I., Horvitz, H.R. & Hirschberg, C.B. SQV-7, a protein involved in Caenorhabditis elegans epithelial invagination and early embryogenesis, transports UDP-glucuronic acid, UDP-N-acetylgalactosamine, and UDP-galactose. Proc Natl Acad Sci U S A 98, 3738–43 (2001).

39. Drew, D. & Boudker, O. Shared Molecular Mechanisms of Membrane Transporters. Annu Rev Biochem 85, 543–72 (2016).

40. Milla, M.E. & Hirschberg, C.B. Reconstitution of Golgi vesicle CMP-sialic acid and adenosine 3’-phosphate 5’-phosphosulfate transport into proteoliposomes. Proc Natl Acad Sci U S A 86, 1786–90 (1989).

41. Weis, W.I. & Kobilka, B.K. The Molecular Basis of G Protein-Coupled Receptor Activation. Annu Rev Biochem 87, 897–919 (2018).

42. Caffaro, C.E. & Hirschberg, C.B. Nucleotide sugar transporters of the Golgi apparatus: from basic science to diseases. Acc Chem Res 39, 805–12 (2006).

43. Al-Saraireh, Y.M. et al. Pharmacological inhibition of polysialyltransferase ST8SiaII modulates tumour cell migration. PLoS One 8, e73366 (2013).

44. Miyazaki, T., Angata, K., Seeberger, P.H., Hindsgaul, O. & Fukuda, M. CMP substitutions preferentially inhibit polysialic acid synthesis. Glycobiology 18, 187–94 (2008).

45. Kean, E.L., Munster-Kuhnel, A.K. & Gerardy-Schahn, R. CMP-sialic acid synthetase of the nucleus. Biochim Biophys Acta 1673, 56–65 (2004).

46. Kota, J., Gilstring, C.F. & Ljungdahl, P.O. Membrane chaperone Shr3 assists in folding amino acid permeases preventing precocious ERAD. J Cell Biol 176, 617–28 (2007).

47. Newstead, S., Kim, H., von Heijne, G., Iwata, S. & Drew, D. High-throughput fluorescent-based optimization of eukaryotic membrane protein overexpression and purification in Saccharomyces cerevisiae. Proc Natl Acad Sci U S A 104, 13936–41 (2007).

48. Sonoda, Y. et al. Benchmarking membrane protein detergent stability for improving throughput of high-resolution X-ray structures. Structure 19, 17–25 (2011).

49. Hattori, M., Hibbs, R.E. & Gouaux, E. A fluorescence-detection size-exclusion chromatography-based thermostability assay for membrane protein precrystallization screening. Structure 20, 1293–9 (2012).

50. Kabsch, W. Xds. Acta Crystallogr D Biol Crystallogr 66, 125–32 (2010).

51. Collaborative Computational Project, N. The CCP4 suite: programs for protein crystallography. Acta Crystallogr D Biol Crystallogr 50, 760–3 (1994).

52. McCoy, A.J. et al. Phaser crystallographic software. J Appl Crystallogr 40, 658–674 (2007).

53. Schroder, G.F., Levitt, M. & Brunger, A.T. Super-resolution biomolecular crystallography with low-resolution data. Nature 464, 1218–22 (2010).

54. Adams, P.D. et al. PHENIX: a comprehensive Pyth syston-basedem for macromolecular structure solution. Acta Crystallogr D Biol Crystallogr 66, 213–21 (2010).

55. Vagin, A.A. et al. REFMAC5 dictionary: organization of prior chemical knowledge and guidelines for its use. Acta Crystallogr D Biol Crystallogr 60, 2184–95 (2004).

56. Emsley, P. & Cowtan, K. Coot: model-building tools for molecular graphics. Acta Crystallogr D Biol Crystallogr 60, 2126–32 (2004).

57. Eckhardt, M., Gotza, B. & Gerardy-Schahn, R. Mutants of the CMP-sialic acid transporter causing the Lec2 phenotype. J Biol Chem 273, 20189–95 (1998).

58. Takeshima-Futagami, T. et al. Amino acid residues important for CMP-sialic acid recognition by the CMP-sialic acid transporter: analysis of the substrate specificity of UDP-galactose/CMP-sialic acid transporter chimeras. Glycobiology 22, 1731–40 (2012).

59. Chan, K.F., Zhang, P. & Song, Z. Identification of essential amino acid residues in the hydrophilic loop regions of the CMP-sialic acid transporter and UDP-galactose transporter. Glycobiology 20, 689–701 (2010).

60. Forrest, L.R. Structural Symmetry in Membrane Proteins. Annu Rev Biophys 44, 311–37 (2015).

61. Vergara-Jaque, A., Fenollar-Ferrer, C., Kaufmann, D. & Forrest, L.R. Repeat-swap homology modeling of secondary active transporters: updated protocol and prediction of elevator-type mechanisms. Front Pharmacol 6, 183 (2015).

