## Supplemental information for "Structural basis for the delivery of activated sialic acid into Golgi for sialyation"

### Supplementary Information

**Table 1. Data collection and refinement statistics.**

| | CST-CMP | CST-*apo* |
|---|---|---|
| **Data Collection** | | |
| Space group | C 1 2 1 | P1 |
| Cell dimensions | | |
| *a, b, c* (Å) | 89.38, 181.00 53.08 | 51.58, 69.46, 102.03 |
| α, β, γ (°) | 90, 90, 90 | 82.83, 83.43, 68.22 |
| Resolution (Å) | 45.79 - 2.8 (2.9 - 2.8) | 47.77 – 3.40 (3.52-3.40) |
| Wavelength (Å) | 0.97625 | 0.97625 |
| $R_{sym}$ or $R_{merge}$ (%)# | 18.85 (194.60) | 12.20 (132.30) # |
| $I / \sigma I$ | 7.59 (0.74) | 7.30 (1.40) |
| Completeness (%) | 99.8 (99.9) | 99.2 (97.7) |
| Redundancy | 7.1 (7.2) | 6.4 (5.1) |
| CC1/2 | 0.88 (0.56) | 0.98 (0.53) |
| **Refinement** | | |
| Resolution (Å) | 45.83 - 2.8 (2.9 - 2.8) | 21.95-3.40 (3.52-3.40) |
| Unique reflections | 19661 (2050) | 16820 (1689) |
| Rwork / Rfree (%) | 23.94 / 25.28 | 26.79 / 28.59 |
| No. atoms | 4897 | 4290 |
| Protein | 4817 | 4290 |
| Ligand/ion | 42 | |
| Water | 38 | |
| *B*-factors | | |
| Protein | 75.34 | 174.71 |
| Ligand/ion | 131.29 | |
| Water | 51.31 | |
| R.m.s deviations | | |
| Bond lengths (Å) | 0.007 | 0.011 |
| Bond angles (°) | 1.03 | 1.49 |
| Ramachandran favoured (%) | 97.55 | 97.66 |
| Ramachandran outliers (%) | 0.00 | 0.00 |

***Values in parentheses are for highest resolution shell.**

**# Three separate datasets were merged together.**

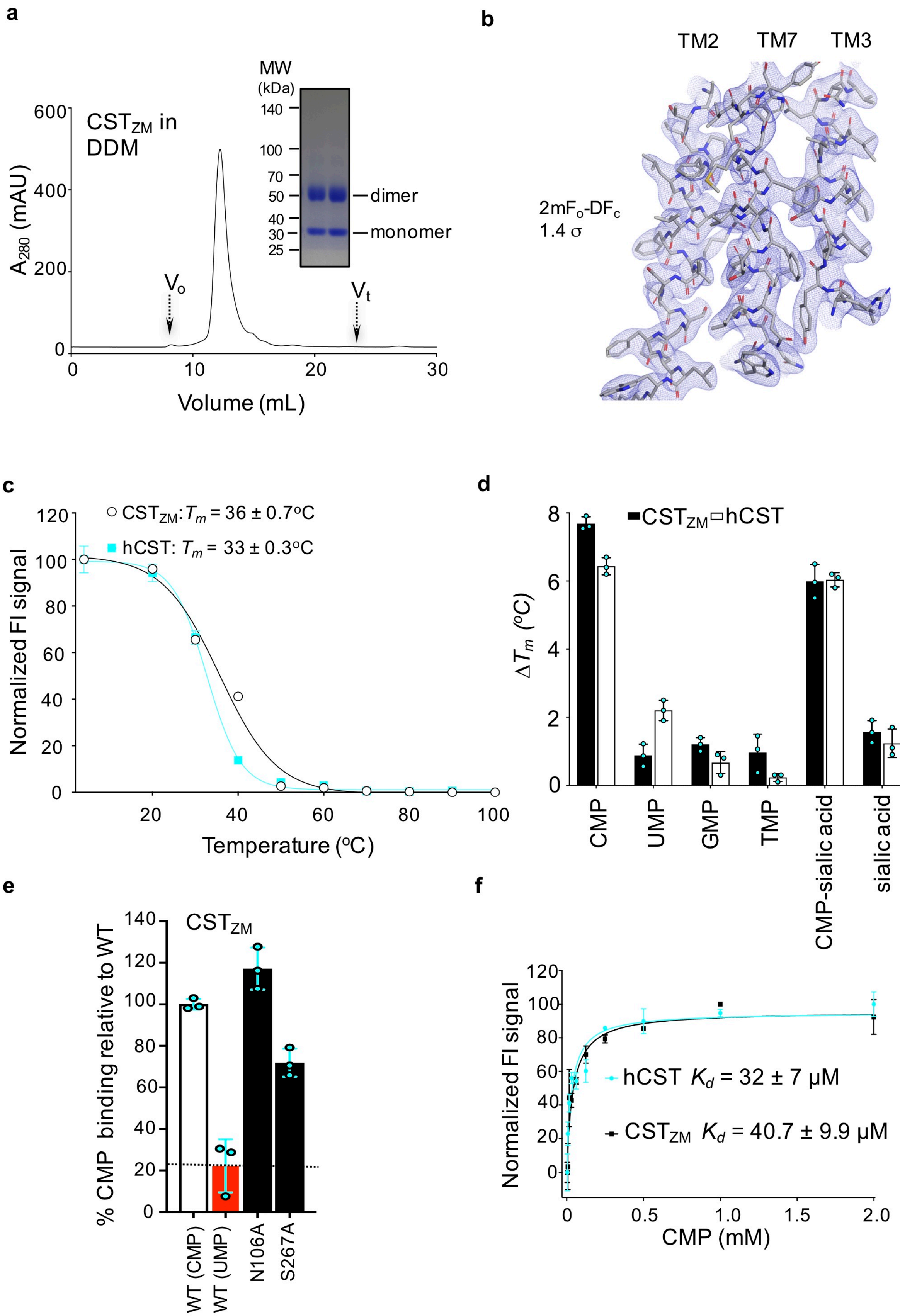
a
b
c
d
e
f
MW (kDa)
140
100
70
50
40
30
25
dimer
monomer
CST$_{ZM}$ in DDM
$A_{280}$ (mAU)
Volume (mL)
$V_o$
$V_t$
TM2
TM7
TM3
2mF$_o$-DF$_c$
1.4 σ
CST$_{ZM}$: $T_m$ = 36 ± 0.7°C
hCST: $T_m$ = 33 ± 0.3°C
Normalized FI signal
Temperature (°C)
CST$_{ZM}$ hCST
$\Delta T_m$ (°C)
CMP
UMP
GMP
TMP
CMP-sialic acid
sialic acid
CST$_{ZM}$
% CMP binding relative to WT
WT (CMP)
WT (UMP)
N106A
S267A
hCST $K_d$ = 32 ± 7 μM
CST$_{ZM}$ $K_d$ = 40.7 ± 9.9 μM
Normalized FI signal
CMP (mM)

**Supplementary Fig. 1| Stability and substrate binding to $CST_{ZM}$ and hCST.** **a,** Size-exclusion chromatogram of $CST_{ZM}$ and corresponding SDS-PAGE-gel with DDM purified $CST_{ZM}$ migrating predominantly as a dimer. **b,** Representative portions of the $2mF_o$-$DF_c$ electron density map (1.4σ), which is shown here for the TM segments involved in CMP binding. **c,** The GFP-TS melting curves using GFP-fusions for $CST_{ZM}$ in crude-detergent solubilized membranes (black open circles) and for hCST (cyan filled squares); errors bars, s.e.m.; n = 3 and the values reported for the apparent $T_m$ are the mean ± s.e.m. of the data fitting. **d,** The apparent $T_m$ for $CST_{ZM}$ and hCST were determined as in A. and further in the presence of 1 mM CMP, 1 mM UMP, 1mM GMP, 1 mM TMP, 10 mM CMP-sialic acid or 10 mM sialic acid. The difference in melting temperature $\Delta T_{\mathrm{m}}$ is shown for $CST_{ZM}$ (filled bars) and hCST (open bars); errors bars are the s.e.m. 3 independent experiments (cyan-filled circles). Markedly, absolute $\Delta T_{\mathrm{m}}$ differences between CMP and UMP are equivalent to that measured using the CPM assay to estimate binding differences between ATP and AMP in the unrelated ADP/ATP exchanger (Majd et al., 2018). **e,** CMP binding to $CST_{ZM}$ using GFP-fusions as estimated by a shift in $\Delta T_{\mathrm{m}}$ after addition of 1 mM CMP to WT (open bars), and to mutants of residues not interacting with crystallographic waters in the cavity, but are present in close proximity to the water network (filled bars). Non-specific binding was estimated with addition of UMP (red bars). **f,** The GFP-TS assay was used to determine binding of CMP to purified GFP-fusions of $CST_{ZM}$ (black squares) and hCST (cyan squares). Binding affinities ($K_{\mathrm{d}}$) were calculated from data points recorded over a range of CMP concentrations, and these were fitted by non-linear regression using data from 3 independent experiments (the values reported are the mean ± s.e.m. of the fit and were equivalent to previously obtained $K_{\mathrm{d}}$ estimates using detergent-solubilised membranes (Nji et al., 2018); see Methods).

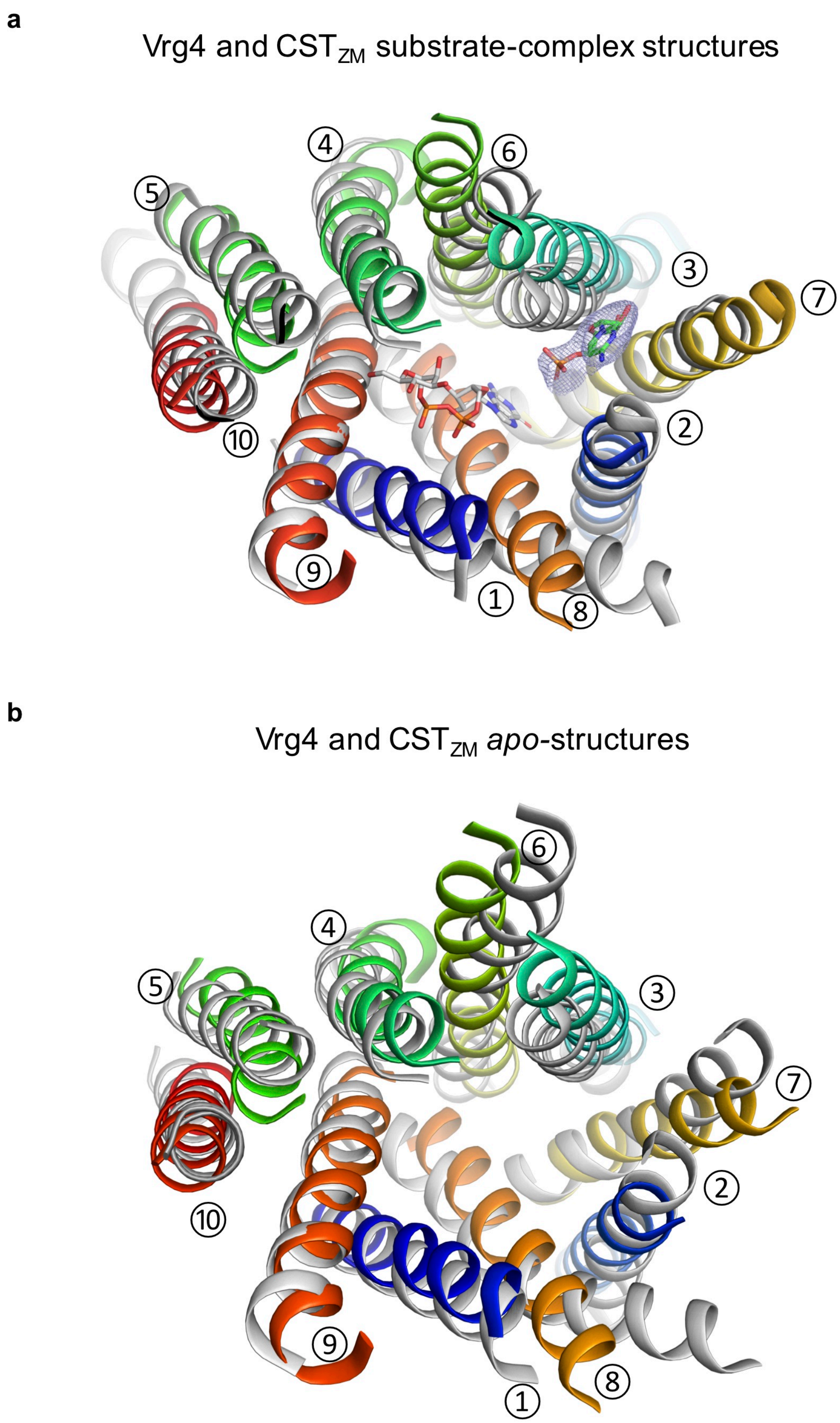


**Supplementary Fig. 2| Superimposition of *apo*- and substrate-bound outward-facing $CST_{ZM}$ and Vrg4 structures. a,** Cartoon representation of outward-facing 2.8Å resolution structure of $CST_{ZM}$ in complex with CMP (colored as in Fig. **1d**) and the 3.6Å outward-facing structure of Vrg4 in complex with GDP-mannose (PDB id: 5OGK; light grey), as viewed from the top of the membrane. The CMP and GDP-mannose moieties from $CST_{ZM}$ and Vrg4 respectively, are shown as sticks. The polder (OMIT) map(Liebschner et al., 2017) for CMP contoured at 3σ is also shown. **b,** As in **a.,** the *apo* 3.4Å outward-facing structure of $CST_{ZM}$ (colored as in Fig. **1d**) and the *apo* 3.2Å resolution outward-facing Vrg4 (PDB id: 50GE; light grey) structures.

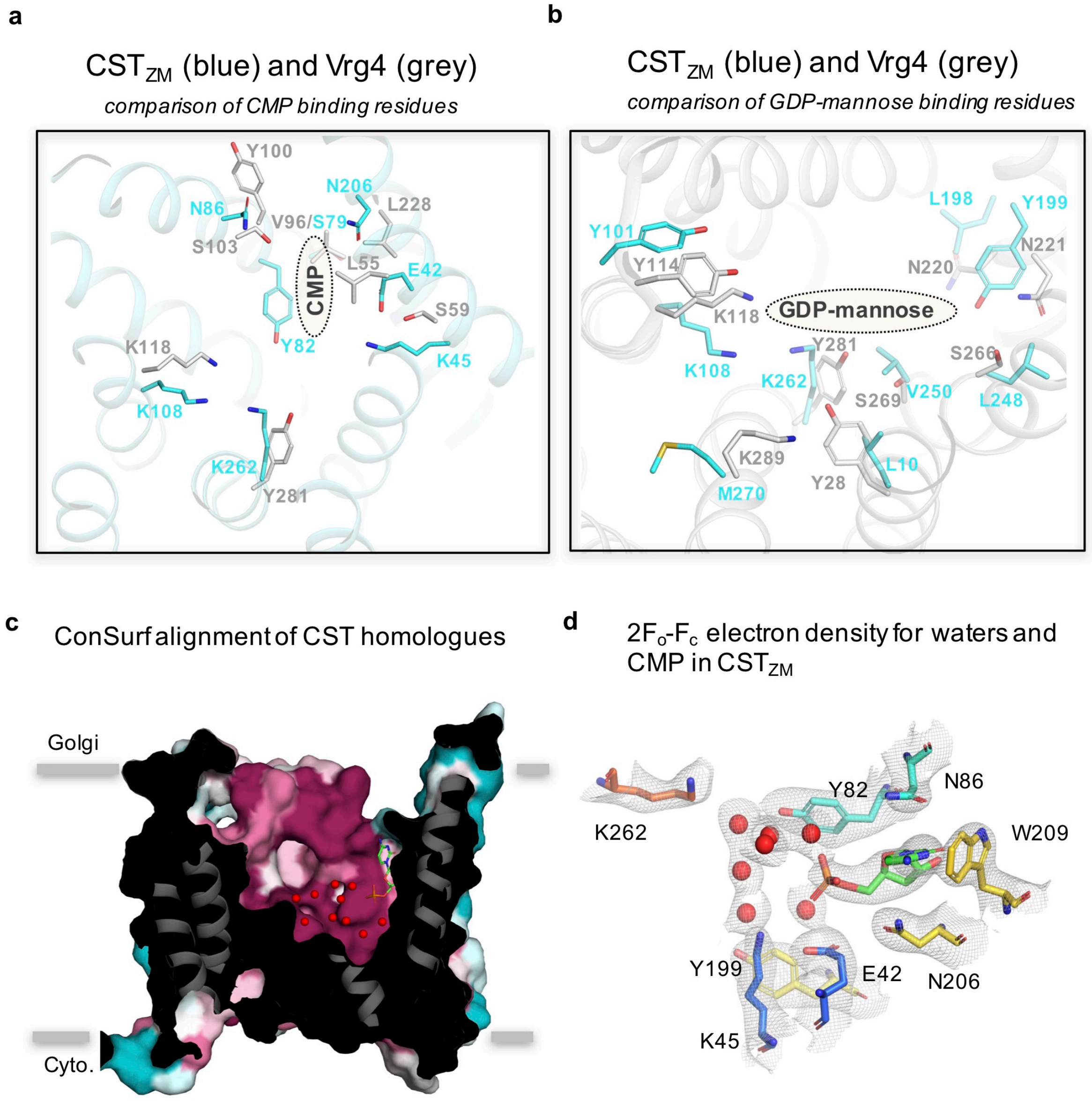


**Supplementary Fig. 3| Comparison of substrate-binding residues in $CST_{ZM}$ and Vrg4 are very different. a,** Cartoon representation illustrating the pronounced differences of the substrate-binding site residues when comparing the outward-facing $CST_{ZM}$ structure in complex with CMP (colored cyan) and the outward-facing Vrg4 structure in complex with GDP-mannose (colored light-grey). The position of the bound CMP is shown as an oval and residues as sticks. **b,** Cartoon representation illustrating the pronounced differences of the substrate-binding site residues in the Vrg4 structure in complex with GDP-mannose (colored grey) when compared to the structure of $CST_{ZM}$ in complex with CMP (colored cyan). The position of the GDP-mannose is shown as an oval and residues as sticks. **c,** Slab through the outward-facing $CST_{ZM}$ as viewed within the plane of membrane with CMP (shown as yellow sticks). All cavity waters in $CST_{ZM}$ are shown as red spheres, which was generated by overlaying separately to the sliced surface. To visualise structural sequence conservation, the ConSurf server was used with default settings (colored based on sequence conservation; low-cyan to high-dark-red)(Landau et al., 2005). The sequence alignment for conservation analysis was generated using Clustal Omega by comparing 63 $CST_{ZM}$ homologues with sequence identity ranging from 23 to 91 %. **d,** Modelled positions and $2mF_O$-$DF_C$ electron density map (1.0 $\sigma$) for the observed waters and CMP in the $CST_{ZM}$ complex and the residues involved in water coordination. Residues and CMP are shown as sticks and waters as red spheres.

**a**

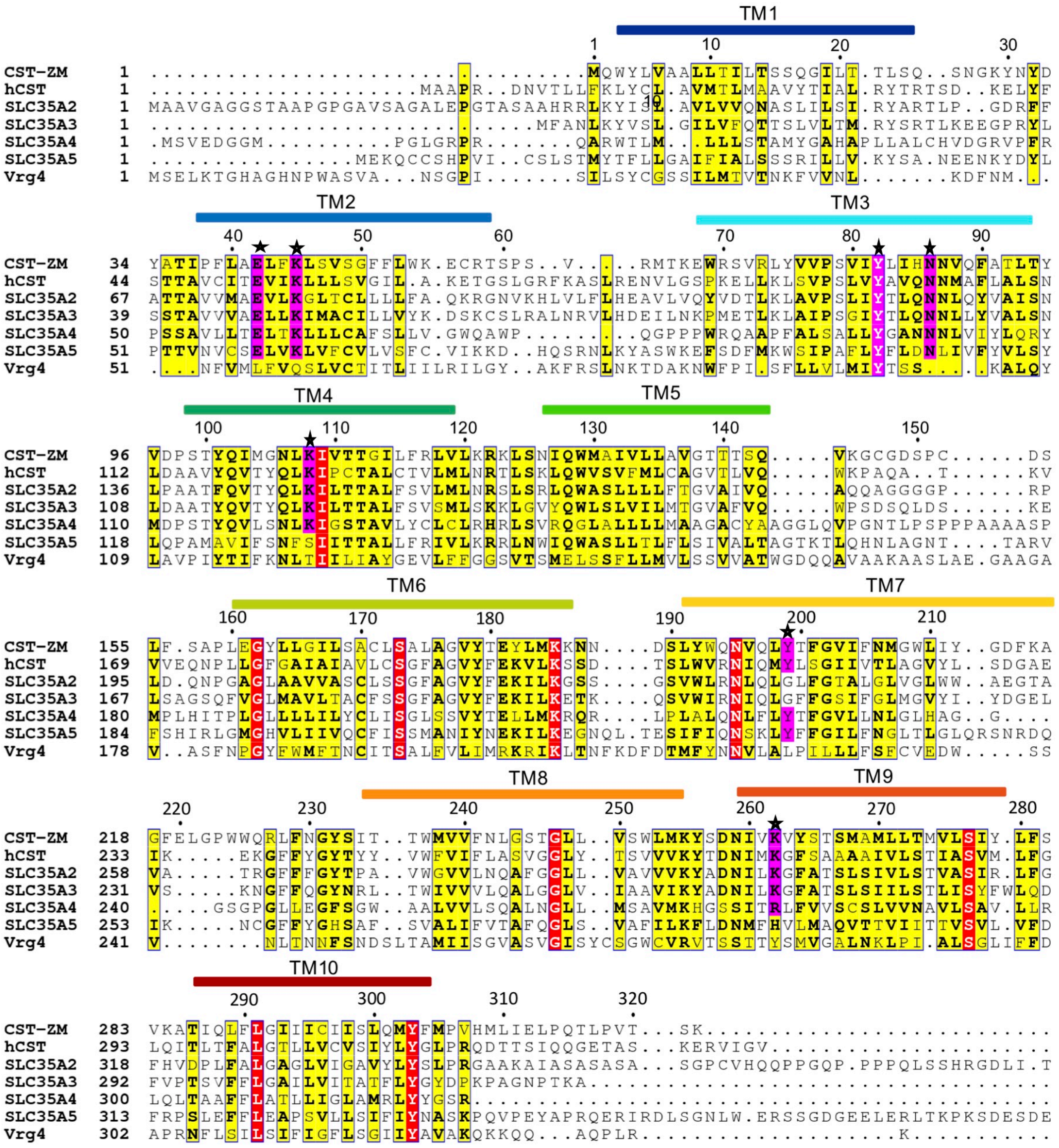


**b**

| | CSTZM | hCST(A1) | SLC35A2 | SLC35A3 | SLC35A4 | SLC35A5 | Vrg4 |
|---|---|---|---|---|---|---|---|
| **CSTZM** | 100.00 | | | | | | |
| **hCST(A1)** | 27.65 | 100.00 | | | | | |
| **SLC35A2** | 27.71 | 43.75 | 100.00 | | | | |
| **SLC35A3** | 26.45 | 39.50 | 53.27 | 100.00 | | | |
| **SLC35A4** | 30.72 | 24.41 | 28.34 | 25.08 | 100.00 | | |
| **SLC35A5** | 25.40 | 24.10 | 25.62 | 26.48 | 21.17 | 100.00 | |
| **Vrg4** | 16.84 | 13.36 | 14.01 | 14.53 | 16.44 | 18.01 | 100.00 |

**Supplementary Fig. 4| Sequence alignment of *Zea mays* CST, *human* CST (SLC35A1), *human* SLC35A2-A5, and *S. cerevisiae* Vrg4. a,** Clustal Omega sequence alignment of selected nucleotide-sugar transporters. Transmembrane helices of *Zea mays* CST are indicated above the alignment and colored in rainbow (as in Fig. **1d**). Strictly conserved residues are highlighted in red boxes and highly conserved residues are highlighted in yellow boxes. Pink boxes, marked by stars, represent conserved residues involved in substrate binding. For sake of clarity, 179-218 loop residues unique to SLC35A5 were not included in the alignment. Notably, Tyr82 in $CST_{ZM}$ is conserved with Tyr100 in Vrg4 at the sequence level, but is not structurally conserved (Supplementary Fig. **3a**). **b,** Table showing the percentage sequence identity of the Clustal Omega alignment shown in **a**.

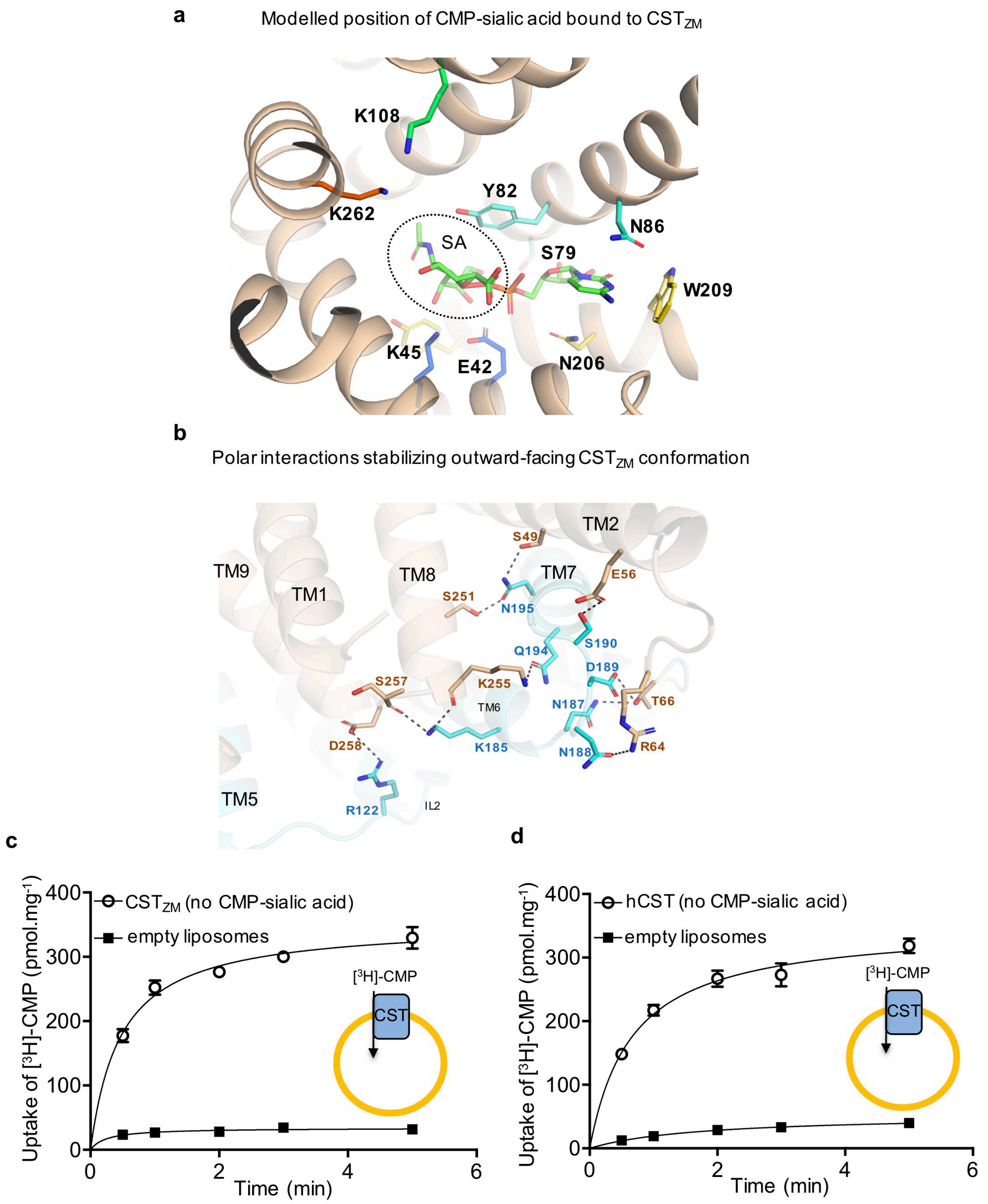


**Supplementary Fig. 5| Modelled position of CMP-sialic acid, inter-domain interactions, and facilitative activities. a,** The substrate-binding site in the outward facing $CST_{ZM}$ structure with modelled CMP-sialic acid (TMs in light brown). The CMP-sialic acid is shown as green sticks, and residues coordinating the modelled CMP-sialic acid as sticks colored in rainbow (as in Fig. **1d**). **b,** Cartoon representation showing the residues which by polar interactions stabilize the two transport bundles (colored cyan and wheat respectively) in the outward-facing conformation. **c,** Time-dependent uptake of [$^3$H]-CMP by $CST_{ZM}$ (open circles). Non-specific uptake was estimated with empty liposomes (filled squares). **d**, Time-dependent uptake of [$^3$H]-CMP by hCST (open circles). Non-specific uptake was estimated with empty liposomes (filled squares). In all experiments errors bars, s.e.m.; n = 3.
